# Loss of LanC-like proteins delays post-injury regeneration of aging skeletal muscles

**DOI:** 10.64898/2026.05.15.725287

**Authors:** Adriana Reyes-Ordoñez, Tianhui Hina Zhou, Tarun C. Rao, Pallob Barai, Wilfred A. van der Donk, Jie Chen

**Affiliations:** Department of Cell & Developmental Biology, Carle Illinois College of Medicine, University of Illinois at Urbana-Champaign, Illinois, USA; Department of Biochemistry, Carle Illinois College of Medicine, University of Illinois at Urbana-Champaign, Illinois, USA; Department of Bioengineering, Carle Illinois College of Medicine, University of Illinois at Urbana-Champaign, Illinois, USA; Department of Chemistry, Carle Illinois College of Medicine, University of Illinois at Urbana-Champaign, Illinois, USA; Howard Hughes Medical Institute, Carle Illinois College of Medicine, University of Illinois at Urbana-Champaign, Illinois, USA; Cancer Center at Illinois, Carle Illinois College of Medicine, University of Illinois at Urbana-Champaign, Illinois, USA; Department of Biomedical and Translational Sciences, Carle Illinois College of Medicine, University of Illinois at Urbana-Champaign, Illinois, USA

**Keywords:** LanCL, muscle regeneration, inflammatory response, aging

## Abstract

The adult skeletal muscle regenerates robustly upon injury, but this regenerative capacity rapidly declines with age. In this study, we identify the lanthionine synthetase C-Like (LanCL) proteins, mammalian homologs of the bacterial peptide cyclase LanC, as positive regulators of muscle regeneration in middle-aged mice. In a barium chloride-induced injury model, we found the protein levels of LanCL1 and LanCL2 to increase during an early phase of regeneration in middle-aged (12-month-old) but not young adult (4-month-old) mice. Utilizing a mouse line lacking all three LanCL proteins (LanCL triple KO or LTKO), we examined a potential role of LanCL in injury-induced muscle regeneration. Consistent with an age-dependent function of LanCL, we observed a delayed regeneration of the tibialis anterior (TA) muscle after injury, as reflected by reduced regenerating myofiber sizes at day 7 after injury (D7AI) in middle-aged (but not young) LTKO compared to age-matched WT mice. The pool size of quiescent satellite cells (Pax7+) was comparable between 12-month-old LTKO and WT muscles without injury. However, the number of Pax7+ cells was significantly higher in regenerating LTKO muscles at both D3AI and D5AI, accompanied by reduced numbers of MyoD+ and MyoG+ cells. The percentage of proliferating Pax7+ cells was also higher in LTKO muscles at D5AI. Furthermore, we detected elevated expression of pro-inflammatory cytokines in regenerating LTKO muscles, while the number of macrophages was similar comparing LTKO and WT muscles. Taken together, our observations suggest that in aging muscles LanCLs are important for proper timing of inflammation resolution and regeneration upon injury.

**New & Noteworthy:** Physiological roles of the mammalian homologs of bacterial LanC, LanCLs, are poorly understood. Our work uncovers a function of LanCLs in post-injury regeneration of aging skeletal muscles. Middle-aged LanCL triple KO mice displayed a delay in satellite cell differentiation and regenerative myofiber formation, as well as persistent inflammatory cytokine expression, suggesting that LanCLs may have an age-dependent role in modulating inflammation in the injured muscles to facilitate regeneration.

## INTRODUCTION

Lanthionine synthetase C-Like protein 1, 2, and 3 (LanCL1, 2, 3) are mammalian homologs of the bacterial peptide cyclase LanC, which is involved in the production of lanthipeptides, members of a natural product family called the ribosomally synthesized and post-translationally modified peptides (1). Lanthipeptides are characterized by lanthionine crosslinks formed from Ser and Cys residues. The Ser is dehydrated to the dehydroamino acid dehydroalanine (Dha), and LanC catalyzes the addition of Cys to the Dha to form the lanthionine crosslink (2). However, LanCLs are not involved in lanthionine synthesis in mammals (3). These evolutionarily conserved proteins were functionally uncharacterized until the early 2000s, but by now multiple independent studies have reported the involvement of LanCL1 and LanCL2 in redox homeostasis (4–11), protein damage control (12), cell survival (5, 7, 13), and immune responses (14–17). Like LanC, LanCLs are zinc-binding proteins with evolutionarily conserved active site residues (18). However, instead of catalyzing intramolecular ring formation through the addition of Cys to Dha, LanCLs catalyze C-glutathionylation of dehydroamino acids generated at protein phosphorylation sites, which may serve to protect the mammalian proteome from reactive electrophiles (12). These findings lead to the speculation that LanCLs may be involved in protein damage control, especially during aging.

Both LanCL1 and LanCL2 are highly expressed in the brain, and LanCL1 has been reported to provide protection against chronic neurodegeneration (7). LanCL1 has also been identified as a critical sensor and regulator of cellular glutathione (GSH) levels through negative regulation of Cystathionine β-Synthase (CBS) (11), making it an antioxidant protein that promotes neuron survival and cancer cell proliferation (5, 7, 8). Such an antioxidant role was also observed for LanCL2 in the regulation of testicular redox homeostasis (8). In addition, LanCL2 was reported to modulate immune response during host-pathogen interactions (14), and has been suggested to be a potential therapeutic target in treating autoimmune and inflammatory diseases (15, 16, 19). Moreover, LanCL2 was found to regulate lipid metabolism by promoting adipogenic differentiation through regulating PPARγ (20), and to promote cell survival by facilitating mammalian target of rapamycin complex 2 (mTORC2) phosphorylation of the Ser/Thr kinase AKT (13).

LanCL1 and LanCL2 are both expressed in skeletal muscles (21). In muscle cells, LanCL1 and LanCL2 are proposed to be receptors for the phytohormone abscisic acid (ABA), which signals through an AMPK/PGC-1α pathway to stimulate glucose uptake and mitochondrial respiration (22, 23). The adult skeletal muscle has a robust capacity to repair and regenerate upon injury, owing to the resident muscle stem cells (MuSCs) that can undergo activation, proliferation and differentiation to form new myofibers. This process requires the coordinated actions of multiple cell types, working in tandem with MuSC in the regenerative niche. For instance, macrophages play critical roles throughout the early stage of regeneration by clearing damaged tissues and promoting MuSC proliferation and differentiation. Redox signaling has long been recognized to play an essential role in skeletal muscle homeostasis and, in more recent years, has also been established as a strategic regulator of cellular and molecular processes required for efficient skeletal muscle regeneration (24). Consistently, antioxidant enzymes are critically involved in muscle regeneration as they induce angiogenesis and reduce fibrosis (25).

Moreover, muscle repair/regeneration places a high demand on proteostasis. LanCL proteins’ involvement in antioxidation, inflammation, and protein damage control potentially put them at the intersection of key stages of muscle repair. Yet, a role of LanCL proteins in muscle regeneration has not been reported. In the current study we made use of a mouse line lacking expression of all three LanCL proteins (3) and observed that LanCL deletion resulted in delayed muscle regeneration upon acute injury. We further identified a dysregulated inflammatory response as a possible cause of the impaired regeneration.

## Materials and Methods

### Animals

All animal experiments in this study followed protocols approved by the Animal Care and Use Committee at the University of Illinois at Urbana-Champaign. Mice with global knockout of LanCL1/2/3 (LTKO) (3) and wild-type (WT) control mice were maintained on an FVB/NJ background (Jackson Laboratory, stock number 001800). Male and female age-matched LTKO and WT mice were used for experiments at 3 to 4 months of age (young), or 12 to 13 months (middle-aged). Mice were housed up to 5 per cage. The cages were connected to an EcoFlo ventilation system (Allentown Inc.) in a specific-pathogen-free animal facility kept at 23 °C. All animals were maintained on a 12 h/12 h light/dark cycle with ad libitum access to water and normal chow.

### Skeletal muscle injury and regeneration

A barium chloride (BaCl_2_) injury-induced muscle regeneration model was used as previously described (26). Briefly, the mouse was anesthetized by isoflurane in an induction chamber (2-3%) and then maintained under anesthesia via a nosecone (1.5-2%) using a precision vaporizer and an active scavenger system. Tibialis anterior (TA) muscles were then injected with 50 μL of 1.2% (w/v) BaCl_2_ (Sigma, Cat. #202738) dissolved in saline (injured) or just saline (uninjured control). At 5, 7, or 14 days after injection, the injected TA muscles were collected from mice euthanized by cervical dislocation under anesthesia. The isolated muscles were snap-frozen in liquid nitrogen-cooled 2-methyl-butane (Fisher Scientific, Cat. #AC126470010) with or without prior embedding in tissue freezing medium TBS (VWR, Cat. #15148-031). The samples were stored at -80 °C until further processing. Samples were excluded from analysis if it was determined that the extent of muscle injury was drastically less than expected (e.g., <30% of the section area containing centrally nucleated myofibers), likely due to mis-injection. To minimize batch and operator effects, every round of injury and tissue collection was performed for both genotypes (WT and LTKO) by the same investigator using the same lot of materials and reagents. Data from both sexes were combined unless otherwise indicated in the figures.

### CSA measurement

Sections of 10-µm thickness of the TA muscle mid-belly were obtained using a cryostat (Microm HM550; ThermoFisher Scientific) at -20 °C and placed on charged slides, followed by staining with Hematoxylin (Sigma-Aldrich, Cat. #GHS316) and Eosin (Sigma-Aldrich, Cat. #HT110116) (H&E). H&E images were acquired using a microscope (DMI 4000B; Leica) with a 20× dry objective (Fluotar, numerical aperture 0.4; Leica) and a camera (RETIGA EXi; QImaging) controlled by the Q-Capture Pro51 software (QImaging). Bright-field images were captured at 12-bit at room temperature (RT) and processed using ImageJ (NIH), where the cross-sectional area (CSA) of centrally nucleated regenerating myofibers was measured. A minimum area of 1,480,000 µm² at the center of the regenerating region of each TA muscle was analyzed for CSA. Investigators were blinded to sample identification while performing CSA quantitation.

### Immunofluorescence imaging and analysis

For immunofluorescence, 10-µm muscle sections were rehydrated with phosphate-buffered saline (PBS), fixed with 3.7% formaldehyde (ThermoFisher Scientific, Cat. #F79-1) for 10 min, permeabilized using 0.1% Triton X-100 (ThermoFisher Scientific, Cat. #BP151) for 5 min, and blocked with 5% BSA (Sigma-Aldrich, Cat. #A7906) and 5% normal goat serum (Jackson ImmunoResearch, Cat. #005-000-001) for 60 min, all at RT. Incubation with primary antibodies was performed overnight at 4 °C. After washing with PBS + 1% BSA, the samples were incubated with Alexa fluor–conjugated secondary antibodies and 1 µg/ml of DAPI (ThermoFisher Scientific, Cat. #P162247) for 60 min at RT. For Pax7 staining, heat-induced epitope retrieval was performed by 5 min pressure-cooking at high pressure (27) after fixation, and samples were cooled down under cold running water before permeabilization. For F4/80 staining, the samples were permeabilized after primary antibody incubation and washing. See Table 1 for detailed information on the antibodies used. Fluorescent images were acquired with the same microscope setup described above, captured as 8-bit images at RT, and processed in ImageJ (NIH). Images were pseudo-colored and adjusted, when necessary, using identical parameters for all samples within the same experiment. A minimum area of 1,088,000 µm² was scored for the number of fluorescent cells. For F4/80 analysis, the corrected total cell fluorescence (CTCF) was quantitated using ImageJ (28).

**Table 1.** Antibodies used in the study.

| Antigen/<br>Antibody species | Source | Catalog # | RRID | Dilution<br>used |
| --- | --- | --- | --- | --- |
| ***Primary antibodies*** |  |  |  |  |
| Pax7/rabbit polyclonal | Aviva Systems Biology | ARP32742_P050 | AB_387565 | 1:400 |
| MyoG/rabbit monoclonal<br>(E9A1S) | Cell Signaling Technology ® | 43098 | AB_3717698 | 1:100 |
| MyoD1/rabbit polyclonal | Proteintech ® | 18943-1-AP | AB_10603467 | 1:500 |
| Ki67/chicken IgY (chimeric),<br>clone SP6 | Abcam | ab320709 | AB_3754680 | 1:100 |
| F4/80/rat monoclonal<br>(clone Cl:A3-1) | BIO-RAD | MCA497 | AB_2098196 | 1:500 |
| LanCL1/mouse monoclonal | Proteintech ® | 68160-1-Ig | AB_2923679 | 1:1000 |
| LanCL2/rabbit polyclonal | Jie Chen Lab (UIUC) | n/a | n/a | 1:1000 |
| GAPDH/rabbit polyclonal | Proteintech ® | 10494-1-AP | AB_2263076 | 1:5000 |
| ***Secondary antibodies*** |  |  |  |  |
| Alexa Fluor 488 AffiniPure<br>F(ab')<sub>2</sub> fragment goat anti-rabbit IgG (H+L) | Jackson ImmunoResearch Inc | 111-546-003 | AB_2338053 | 1:500 |
| Alexa Fluor 488-conjugated<br>donkey anti-rat IgG | Jackson ImmunoResearch Inc | 712-545-150 | AB_2340683 | 1:500 |
| Alexa Fluor 594 AffiniPure<br>F(ab')<sub>2</sub> fragment goat anti-rabbit IgG (H+L) | Jackson ImmunoResearch Inc | 111-586-003 | AB_2338066 | 1:500 |
| Alexa Fluor 488 polyclonal<br>IgG goat anti-chicken IgY<br>(H+L) | Abcam | ab150169 | AB_2636803 | 1:500 |
| Peroxidase AffiniPure goat<br>anti-rabbit IgG (H+L) | Jackson ImmunoResearch Inc | 111-035-003 | AB_2313567 | 1:4000 |
| Peroxidase AffiniPure goat<br>anti-mouse IgG (H+L) | Jackson ImmunoResearch Inc | 115-035-003 | AB_10015289 | 1:4000 |

### mRNA extraction and quantitative real-time PCR

Ice-cold TRI reagent (Sigma-Aldrich, Cat. #T9424) was added to frozen muscles and homogenized using a hand-held homogenizer (ThermoFisher Scientific), followed by total mRNA extraction according to the manufacturer recommendations. mRNA concentrations were measured using a NanoDrop ND-2000C (ThermoFisher Scientific) and 1 μg of the total mRNA was reverse-transcribed using qScript cDNA Synthesis Kit (Quanta Bioscience, Cat. # 95048). Quantitative polymerase chain reaction (qPCR) was performed with SYBR green using a StepOnePlus Real-Time PCR System (Applied Biosystems). The qPCR reactions were run in triplicate for each sample. Samples for comparison were always run on the same qPCR plate.

Investigators were blinded to sample identification when performing the reactions. All the qPCR primers (Table 2) were validated following recommendations by Bustin & Huggett (29). Data were processed using the 2^−ΔΔCt^ method (30) and normalized to the control group (WT). Hprt was reported to be a reliable reference gene for studying gene expression in muscle injury (31), and its levels did not differ when comparing TA muscles of WT and LTKO mice.

**Table 2.**
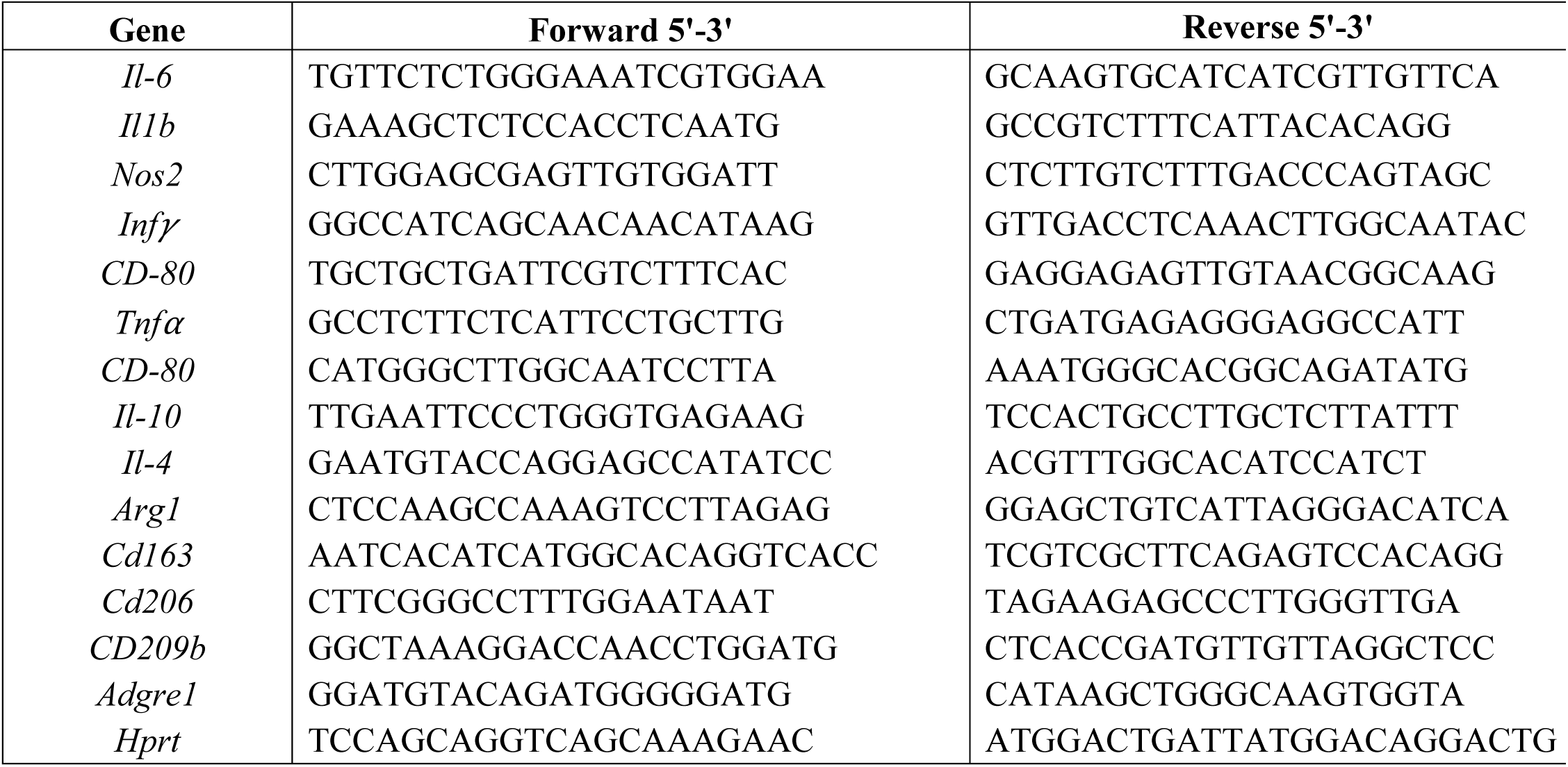
Primers used for qPCR analysis in the study.

### Protein extraction and western blotting

Frozen muscles were mechanically disrupted using a hand-held homogenizer (ThermoFisher Scientific) in ice-cold lysis buffer containing 20 mM Tris-HCl (pH 7.4), 25 mM NaF, 0.1 mM Na_3_VO_4_, 25 mM β-glycerophosphate, 2 mM EDTA, 2 mM EGTA, 0.3% Triton X-100 and 1x protease inhibitor cocktail (Sigma-Aldrich, Cat. #P8340). The homogenates were pre-cleared by centrifugation at 16,200xg for 10 min at 4 °C and protein concentrations were determined using Bradford protein dye (BioRad, Cat. #5000006). Protein samples (15 μg per sample) were separated by sodium dodecyl sulfate-polyacrylamide gel electrophoresis (SDS-PAGE) and transferred to polyvinylidene fluoride (PVDF) membranes (ThermoFisher Scientific, Cat. #PI88520). After blocking with 5% non-fat dry milk in PBS containing 0.1% (v/v) Tween 20 (PBST), membranes were incubated with the primary antibody overnight at 4 °C, followed by washing with PBST and incubation with horseradish peroxidase-conjugated secondary antibody for 30 min at RT. See Table 1 for detailed information on the antibodies used. Chemiluminescent signals were developed using SuperSignal Western Pico PLUS Chemiluminescent Substrate (ThermoFisher Scientific, Cat. #34580) and captured on an iBright CL1000 imaging system (ThermoFisher Scientific). Western blot images were quantitated by densitometry in ImageJ (NIH). Relative protein levels were normalized to GAPDH as the loading control.

### Single-cell RNA sequencing data analysis

We analyzed target gene expression using publicly available mouse skeletal muscle single-cell RNA sequencing (scRNA-seq) datasets. Raw scRNA-seq data from multiple published datasets were obtained, covering early regeneration time points (0, 2, 5, and 7 days after injury) from both young adult (4-7 months) and old (20 months) mice (GEO accession: GSE143437, GSE159500, GSE162172) (32, 33). Processing and analysis of the raw sequencing datasets were performed as previously described (34). Briefly, after quality control, normalization, dimensionality reduction, integration, clustering, and cell type annotation, the expression of Lancl1 and Lancl2 were quantified across defined cell populations and over regeneration time points. Pseudo-bulk differential expression analysis was performed on scRNA-seq data by aggregating raw gene expression levels using the AggregateExpression function in Seurat across cells within each cell type for each biological replicate. Differential expression analysis between different regeneration timepoints was subsequently conducted using DESeq2 on the aggregated counts.

### Statistical analysis

All statistical analyses were performed using R (version 4.5.2; RRID:SCR_001905) within the RStudio integrated development environment (Version 2026.08.2+200; RRID:SCR_000432). Prior to any comparison, data were screened using the Grubbs’ test (two-sided; grubbs.test, R package outliers v0.15) and outliers were eliminated. Sample size (number of animals) at the start of each experiment was determined based on the results and power analysis of the same type of experiments previously performed in our laboratory (e.g., (35, 36)). The exact sample size for each experiment is described in figure legends. All quantitative data are presented as mean ± SD, with individual data points shown. For two-group comparison of means, normality and variance were confirmed using the Shapiro-Wilk and Levene’s test, respectively, followed by an unpaired two-tailed Student’s t-test. For comparison of data from more than two groups, one-way ANOVA was performed, followed by Dunnett’s post-hoc test against the control group. For comparison of myofiber CSA size distribution, the data was analyzed using a linear mixed-effects model (genotype vs size bin interaction with the animal being the random effect). Significant interactions were followed by Fisher’s LSD post-hoc comparison at each bin. Bins of 100 µm^2^ and 200 µm^2^ were applied to D7AI and D14AI, respectively. A difference was considered significant when p < 0.05. All p values of significance are shown directly in the figures.

### Data availability

All data generated or analyzed during this study are included in the manuscript.

### Code availability

All codes used in this study are available on Zenodo: https://zenodo.org/records/22738941

## RESULTS

### Expression of LanCL1 and LanCL2 proteins in regenerating skeletal muscles

LanCL1 and LanCL2 are broadly expressed in most mammalian tissues including the skeletal muscle, whereas LanCL3 expression is very low at the transcript level in all tissues and information on its protein expression is scarce due to the lack of optimal reagents (The Human Protein Atlas (21)). To probe a potential involvement of LanCL1 and LanCL2 in injury-induced muscle regeneration, we started by examining the expression of these proteins in regenerating muscles of wild type mice post-injury. The tibialis anterior (TA) muscle was injured by barium chloride (BaCl_2_) injection and isolated on day 5 or 7 after injury (D5AI or D7AI), followed by protein extraction and western blotting analysis. Uninjured muscles were also collected for analysis (D0). In young adult mice at 3-4 months of age, the expression levels of LanCL1 and LanCL2 did not change markedly comparing regenerating muscles at D5AI or D7AI to uninjured muscles (D0) (Fig. 1A). However, at 12 months of age (middle-aged), expression of both LanCL1 and LanCL2 increased upon injury, at both D5AI and D7AI (Fig. 1B). The mRNA levels of these two genes did not correlate with their protein levels, as Lancl1 mRNA decreased temporarily at D3AI whereas Lancl2 mRNA decreased at D5 and D7AI (Fig. 1C). Hence, LanCL1 and LanCL2 expression during muscle regeneration may be under post-transcriptional regulation. When comparing young and middle-aged muscles without injury, we observed no difference in the protein expression level of LanCL1 or LanCL2 (Fig. 1D). Taken together, our observations imply that LanCL1/2 may have a unique function in injury response of aging muscles.

**Fig. 1.**
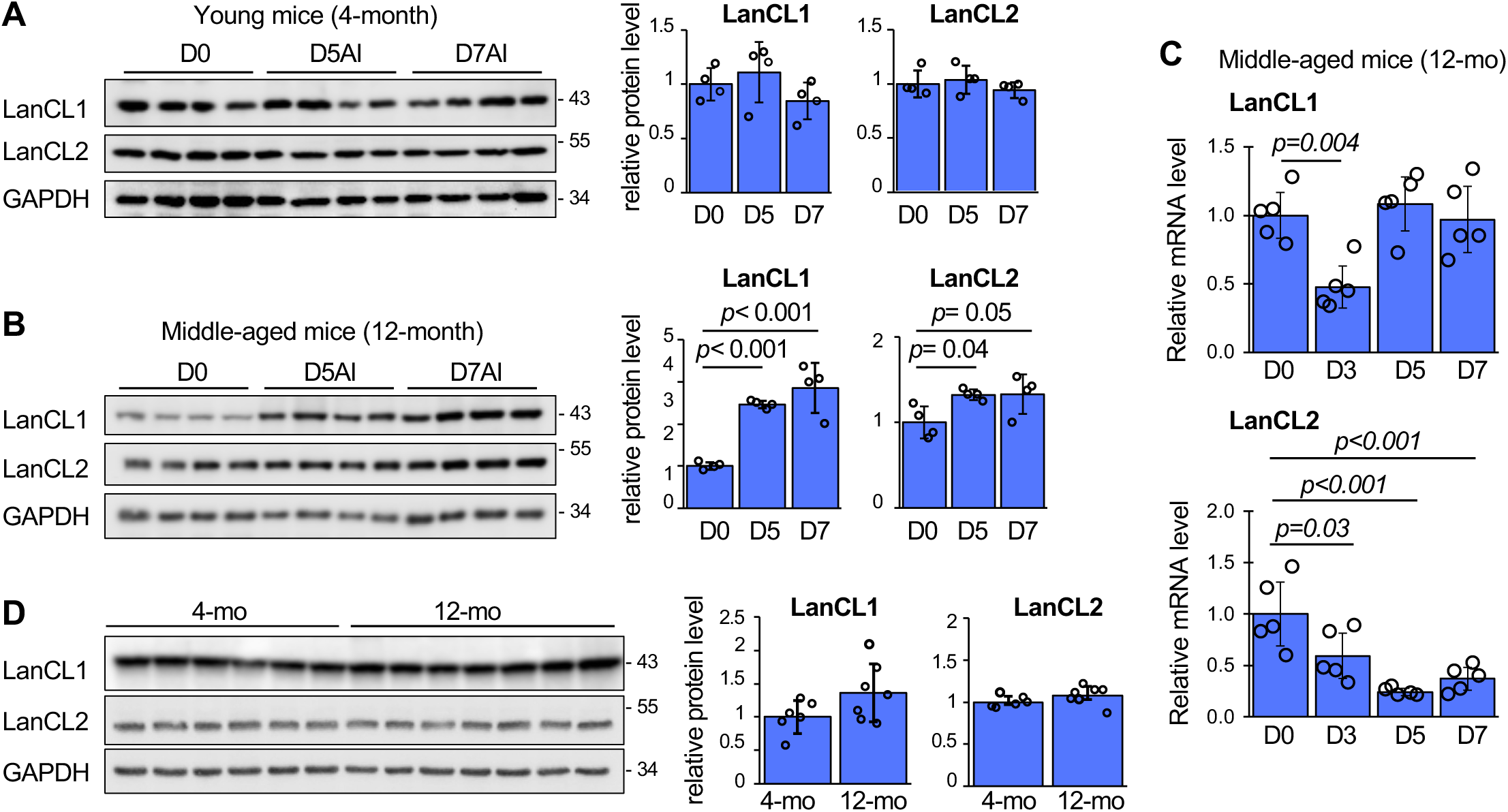
LanCL1 and LanCL2 protein expression during muscle regeneration. (**A-B**) TA muscles of 4-month-old (A) or 12-month-old (B) mice were injected with BaCl_2_. At D5AI and D7AI the injected muscles were isolated and subjected to homogenization and analysis by western blotting. Saline injection served as D0 control. Each experimental group contained 4 animals. Western blots were quantified by densitometry, and GAPDH was used as a reference for relative protein expression levels. Quantification data are presented as mean ± SEM, with individual data points shown. N=4 animals for each condition. (**C**) Mice were treated as described in A-B, and TA muscles isolated at D3, D5, and D7AI were subjected to RNA extraction and qRT-PCR analysis. N=5 animals for each condition. (**D**) TA muscles of 4-month-old or 12-month-old mice without injury were isolated and analyzed by western blotting as described above. N=6-7 animals for each condition. All quantification data are presented as mean ± SD, with individual data points shown. One-way ANOVA was performed to analyze statistical significance of data comparison. The exact p values are indicated on the graphs whenever p < 0.05.

Next, we interrogated the vast single-cell RNA sequencing (scRNA-seq) datasets publicly available (37) to gain cell type-specific insights into the mRNA expression of Lancl genes in regenerating muscles. We performed integration of scRNA-seq data for uninjured (D0) and regenerating TA muscles at D5AI and D7AI for young (4-7 months) and aged (20 months) mice, and analyzed the transcripts of Lancl1 and Lancl2 in the integrated datasets. As expected, Lancl3 mRNA was barely detectable in this dataset. As shown in Fig. 2A, both Lancl1 and Lancl2 genes were widely expressed across all cell types in muscles when combining data from all time points of regeneration. In both young and aged muscles, muscle stem cells (MuSCs) displayed increased levels of Lancl1 mRNA expression on D5AI and D7AI compared to D0 (Fig. 2B). Lancl2 mRNA levels also appeared to increase during regeneration of both young and aged muscles, albeit to lesser degrees (Fig. 2C). Both Lancl1 and Lancl2 expression, in both young and old muscles, were relatively modest in macrophages, the major type of immune cells involved in muscle regeneration (Fig. 2B-C). While the single-cell analysis revealed trends of expression, conclusions about expression level change cannot be drawn from this type of analysis. To gain insights into gene expression across biological replicates from the existing scRNA-seq datasets, we performed pseudo-bulk gene expression analysis for MuSCs. When comparing D5AI and D7AI to D0, no statistically significant change was found for Lancl1 or Lancl2 in aged MuSCs (Fig. 2D). There was also no significant difference in Lancl1 or Lancl2 expression comparing young and aged MuSCs at D5AI or D7AI (Fig. 2D). In conclusion, the mRNA expression patterns derived from scRNA-seq data are distinct from those of protein expression (Fig. 1), suggesting that LanCL1 and LanCL2 expression during muscle regeneration may be under post-transcriptional regulation.

**Fig. 2.**
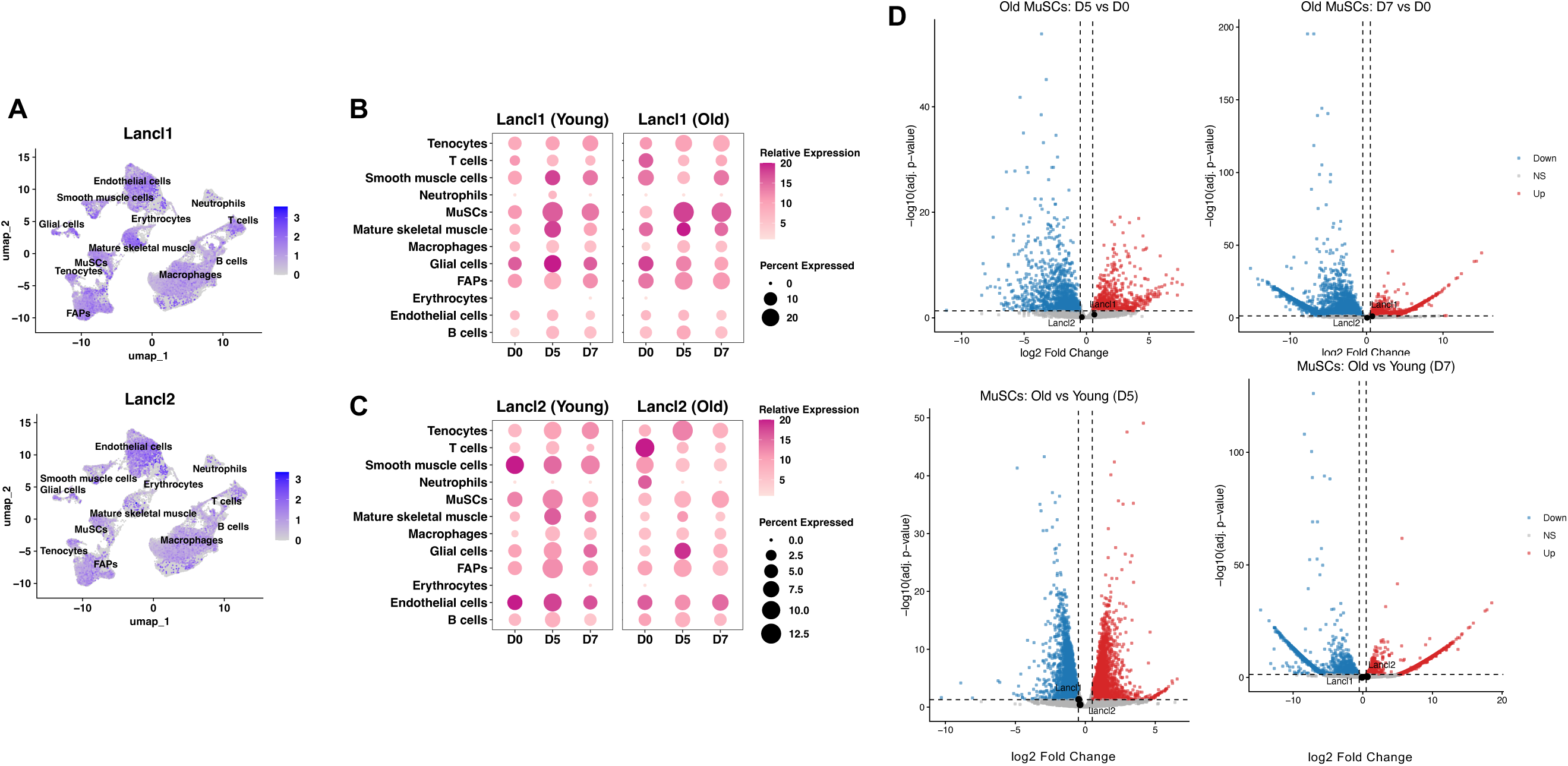
ScRNA-seq analysis of expression of *Lancl1* and *Lancl2* during muscle regeneration. scRNA-seq datasets of mouse TA muscles at D0, D2, D5, and D7AI were integrated and analyzed. (**A**) Single-cell atlases of 170,088 cells combining the timepoints are shown. Data are presented in UMAP to visualize the transcript levels of Lancl1 and Lancl2 in various cell clusters. (**B**) Dot-plots show percentage of cells expressing Lancl1 and relative expression level in various cell clusters of young (4-7 months) and old (20 months) mouse TA muscle at D0, D5, and D7AI. (**C**) Similar to (B) for expression of Lancl2. (**D**) Pseudo-bulk analysis was performed with the scRNA-seq data to identify global transcriptional changes in young and old MuSCs during regeneration. Volcano plots show differentially expressed genes comparing D5AI or D7AI with D0 in old MuSCs (upper) or comparing old and young MuSCs at D5 and D7AI (lower). The positions of Lancl1 and Lancl2 genes are marked on the graphs; neither gene was differentially expressed.

### Loss of LanCL proteins leads to delayed muscle regeneration

The protein expression patterns of LanCL1 and LanCL2 described above implicated these proteins in a potential function during muscle repair in aging. To investigate the role of LanCL proteins, we took advantage of a LanCL triple knockout mouse line with all three Lancl genes systemically disrupted and protein expression abolished (henceforth referred to as LTKO) (3). The LTKO mice do not display any gross phenotype (3). BaCl_2_ injection was performed to injure TA muscles, followed by isolation of the injured muscles at D7AI and D14AI, and quantification of the average cross-sectional area (CSA) of regenerating myofibers to assess the degree of regeneration. In young adult mice, the average regenerating myofiber CSA was not significantly different comparing WT and LTKO muscles at D7AI or D14AI in either male (Fig. 3A) or female mice (Fig. 3B). Although the myofiber size distribution at D14AI in both male and female mice displayed a slight left shift for LTKO compared to WT, neither reached statistical significance. At 12 months of age, LTKO animals exhibited reduced average CSA compared with WT at D7AI and a left shift of the CSA size distribution, observed in both male (Fig. 4A) and female (Fig. 4B) mice, both statistically significant. Those differences were no longer found by D14AI (Fig. 4A&B), although exploratory post-hoc Fisher LSD bin-wise comparisons identified a narrower size distribution in LTKO of both sexes at D14AI. These observations indicate that loss of the LanCL proteins resulted in an age-dependent delay in muscle regeneration. This transient impairment observed in older LTKO mice suggests a possible role for the LanCL proteins in an early stage of regeneration, such as satellite cell activation or differentiation, or inflammatory responses.

**Fig. 3.**
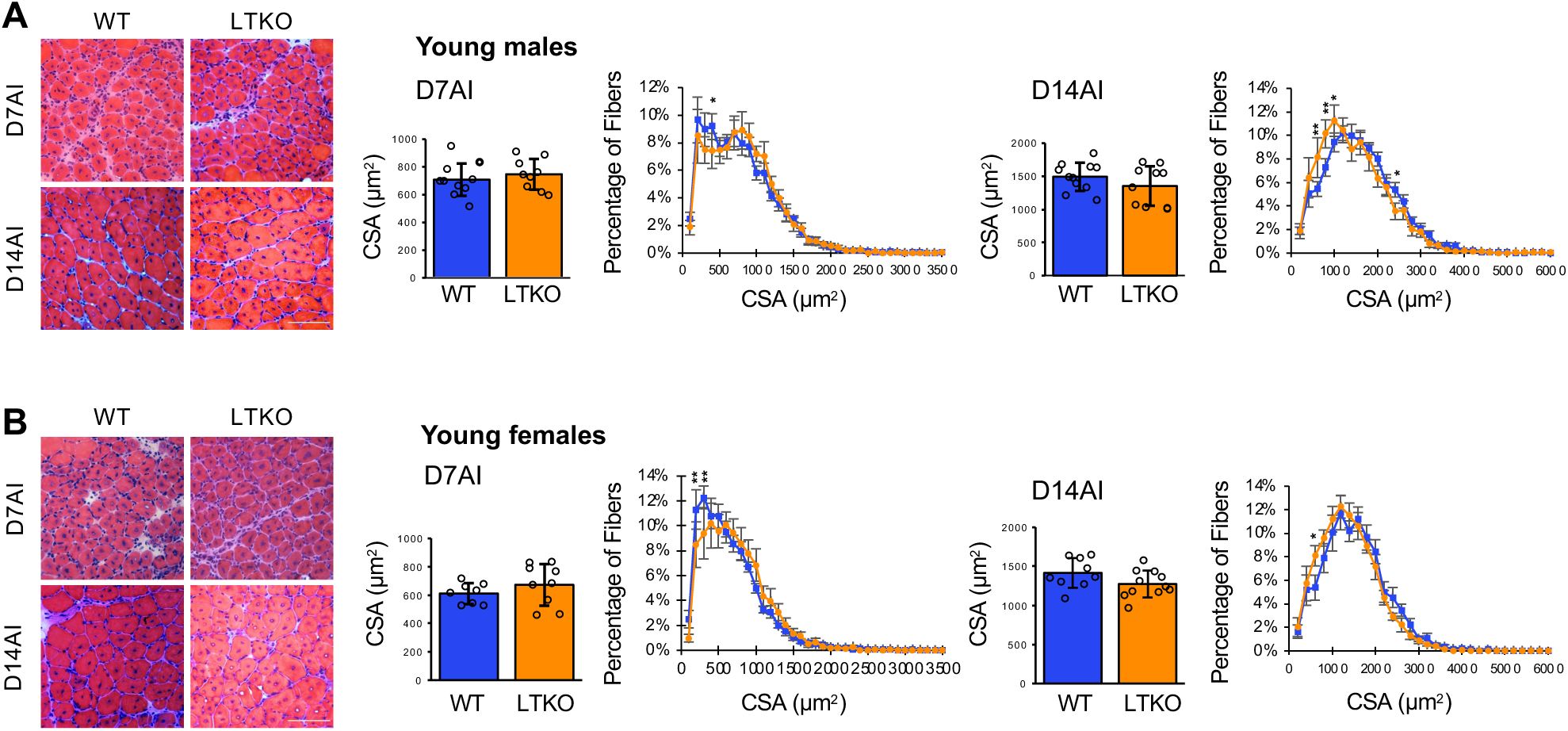
Normal muscle regeneration in young adult LTKO mice. TA muscles of male (**A**) and female (**B**) 4-month-old WT and LTKO mice were subjected to BaCl_2_ injury and isolated at D7AI or D14AI. Cryosections of regenerating muscles were stained with H&E, and cross-sectional area (CSA) of centrally nucleated myofibers was measured. Representative H&E images are shown. Scale bars: 100 µm. Myofiber CSA average and distribution are shown as mean ± SD. For A, N=9-11 per group; for B, N=8-11 per group. Data in blue: WT; orange: LTKO. Two-tailed Student’s t test was performed to analyze statistical significance of any difference in CSA, and no significance was found. Fiber size distribution differences were compared using a linear mixed model two-way ANOVA. No significant difference in the global size was found. An exploratory Fisher’s LSD post-hoc comparison between individual bins was performed, and significant differences were indicated by asterisks: *p < 0.05, **p < 0.01.

**Fig. 4.**
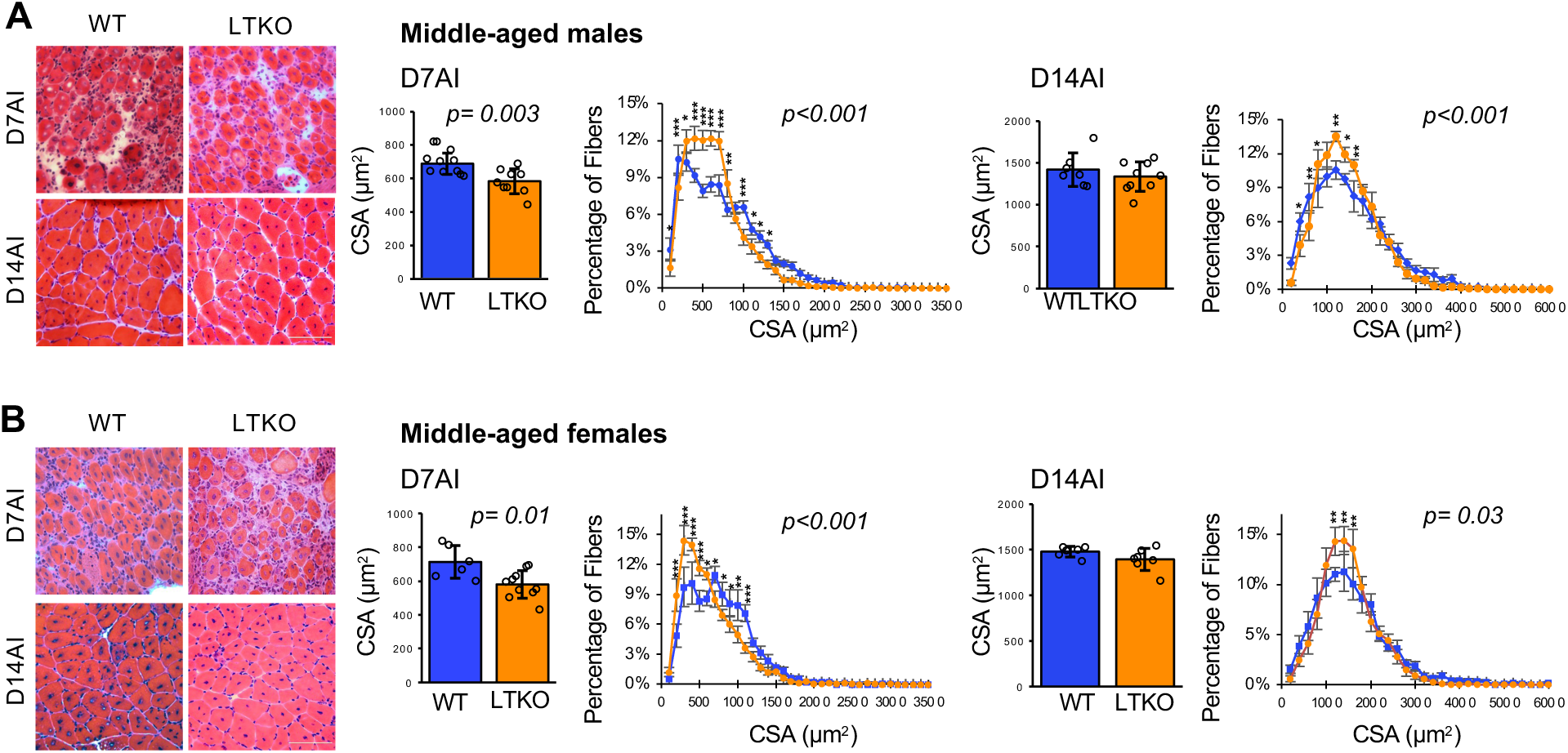
Delayed muscle regeneration in middle-aged LTKO mice. TA muscles of male (**A**) and female (**B**) 12-month-old WT and LTKO mice were subjected to BaCl_2_ injury and isolated at D7AI or D14AI. Cryosections of regenerating muscles were stained with H&E, and cross-sectional area (CSA) of centrally nucleated myofibers was measured. Representative H&E images are shown. Scale bars: 100 µm. Myofiber CSA average and distribution are shown as mean ± SD. For A, N=7-11 per group; for B, N=6-10 per group. Data in blue: WT; orange: LTKO. Two-tailed Student’s t test was performed to analyze statistical significance of differences in CSA, and significant p values are indicated on the graphs. Fiber size distribution differences were compared using a linear mixed model two-way ANOVA, and significant p values are indicated on the graphs. A Fisher’s LSD post-hoc comparison between individual bins was performed, and significant differences are indicated by asterisks: *p < 0.05, **p < 0.01, ***p < 0.001. Post-hoc Fisher LSD bin-wise comparison for overall distribution was also performed, and significant p values can be found on the graphs.

### Satellite cell differentiation is delayed in LTKO muscles

To investigate whether satellite cell behavior was affected by loss of LanCL proteins in 12-month-old muscles, we performed immunohistochemical analysis of TA muscle cross sections before injury (D0) and during regeneration at D3AI and D5AI. Pax7 is known to be expressed in quiescent as well as activated satellite cells, but not in differentiating myocytes. As shown in Fig. 5A the number of Pax7+ cells per cross sectional area was comparable between WT and LTKO muscles prior to injury, suggesting that loss of LanCL proteins did not affect the quiescent muscle stem cell pool. However, during the early phase of regeneration, at both D3AI and D5AI, we observed a markedly higher number of total Pax7+ cells as well as proliferating Pax7+ cells marked by Ki67 in LTKO muscles; at D5AI the percentage of proliferating cells in the Pax7+ pool was also significantly higher in this group (Fig. 5B). Compared to the WT, LTKO muscles contained significantly lower numbers of MyoD+ cells at both D3AI and D5AI (Fig. 5C). Cells expressing myogenin (MyoG), a marker of myogenic differentiation, also decreased in number drastically at D5AI in LTKO muscles (Fig. 5D). Taken together, these observations suggest that LTKO satellite cells may be stalled in the proliferation stage and delayed in their entry into differentiation, which is consistent with the observed delay in new myofiber formation (Fig. 4).

**Fig. 5.**
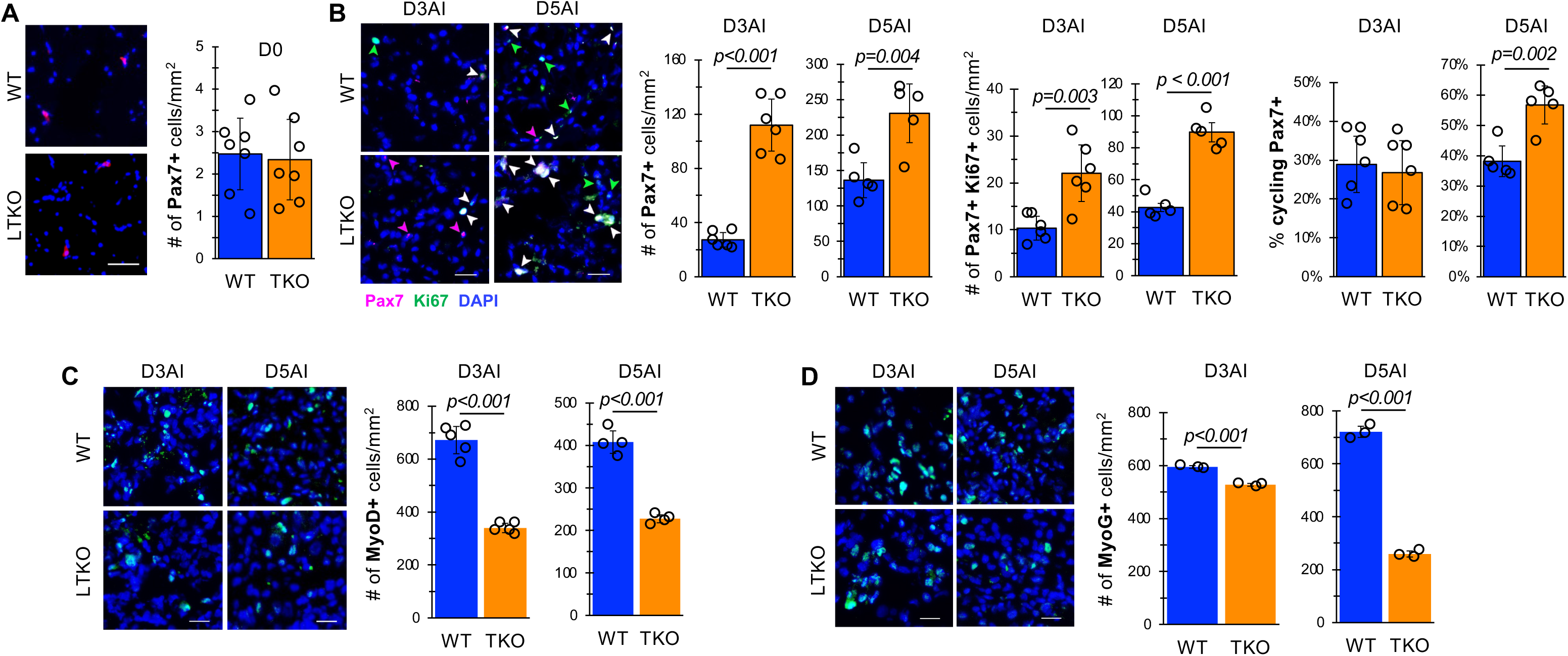
Delayed satellite cell differentiation in regenerating muscles of middle-aged LTKO mice. (A) Uninjured TA muscles of 12-month-old WT and LTKO mice were isolated (D0). Cryosections of muscles were immunostained for Pax7 (magenta) with nuclear staining by DAPI (blue), and Pax7-positive cells were counted. (**B-D**) TA muscles of 12-month-old WT and LTKO mice injured by BaCl_2_ were isolated at D3 or D5AI and subjected to immunostaining for Pax7 (magenta) together with Ki67 (green) (B), MyoD (green) (C), or MyoG (green) (D), with nuclear staining by DAPI (blue). Representative images are shown. Arrowheads indicate Pax7+ only (magenta), Ki67+ only (green), or Pax7+/Ki67+ (white). Scale bars: 20 µm. Quantification data are presented as average number of positive cells per field ± SD, with individual data points shown. N=7 (A), 5-6 (B), 4-5 (C), 3 (D). Blue bars: WT; orange bars: LTKO. Two-tailed Student’s t test was performed to analyze statistical significance for comparison between WT and LTKO at each time point. The exact p values are indicated on the graphs whenever p < 0.05.

### Loss of LanCL proteins delays resolution of inflammation during regeneration

The initial pro-inflammatory response to injury as well as timely resolution of inflammation is critical for injury-induced muscle regeneration. Following acute skeletal muscle injury, infiltrating immune cells, mainly macrophages, adopt an initial pro-inflammatory, debris-clearing, and pro-proliferative program in the first 1-2 days post-injury. After that, the macrophages undergo a transition into a reparative role aiming to support MuSCs differentiation, growth and remodeling. This transition is reported to begin around day 2 and be completed around day 4-5 post injury, following which macrophages decline in number but play a supportive role in myoblast differentiation and fusion as well as contributing to vascular and extracellular matrix remodeling (38–40). To understand how the loss of LanCL proteins could lead to delayed myogenic differentiation and regeneration in aging mice, we set out to investigate the inflammatory response and the shift of gene expression programs from pro-inflammation to pro-regeneration in WT versus LTKO muscles. We performed qRT-PCR to quantitate the mRNA levels of multiple pro- and anti-inflammatory markers in injured TA muscle at D5AI, a key time point in the inflammatory-to-reparative transition.

We examined the expression of the following mRNAs: markers associated with pro-inflammatory activity (M1-like) macrophage activation (Il6, Il1b, Nos2, Tnfa, Ifng, Cd80 and Cd86), markers typically associated with anti-inflammatory transition (Il10 and Il4), and markers associated with reparative (M2-like) macrophages (Arg1, Cd163, Cd206 and Cd209b).

Comparing 12-month-old LTKO and WT muscles, we observed significantly elevated levels of the proinflammatory cytokines Il6, Il1b, Ifng, Tnfα, and Nos2 at D5AI (Fig. 6A), a time that anti-inflammatory response should be taking place (38–40). This observation suggests that LTKO muscles were in a prolonged pro-inflammatory state, which might have been responsible for the delayed satellite cell differentiation. No difference was found in the expression of Adgre1, a pan-macrophage marker, between WT and LTKO at either D3AI or D5AI (Fig. 6B). The number of F4/80+ cells (all macrophages) in TA muscles was also comparable between the two genotypes at both time points (Fig. 6C).

**Fig. 6.**
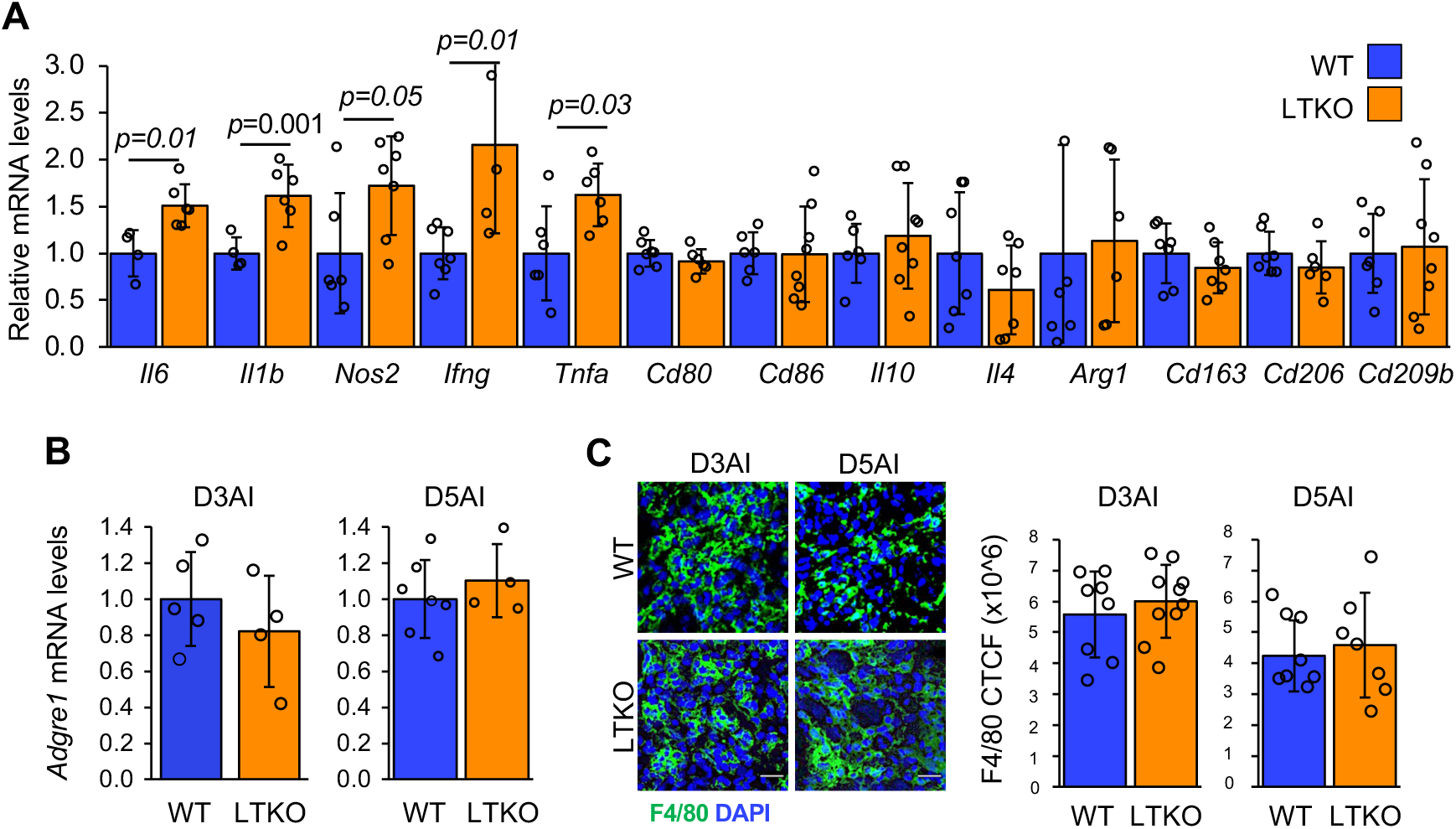
Delayed resolution of inflammation in LTKO muscles after injury. (**A-B**) TA muscles of 12-month-old WT (N=4-7) and LTKO (N=4-8) mice were subjected to BaCl_2_ injury and isolated at D5AI (A) or D3AI and D5AI (B), followed by RNA extraction and qPCR analysis of relative levels of mRNA expression. (**C**) TA muscles treated as in (A-B) were cryosectioned and immunostained with anti-F4/80 (green) with nuclear staining by DAPI (blue). For WT, N=8 for both D3AI and D5AI; for LTKO, N=10 for D3AI and N=7 for D5AI. Representative images are shown. Scale bars: 20 µm. F4/80 fluorescent signals were quantified as corrected total cell fluorescence (CTCF). Quantitative data are presented as mean ± SD with individual data points shown. Two-tailed Student’s t test was performed to analyze statistical significance. The exact p values are indicated on the graphs whenever p < 0.05.

## DISCUSSION

In this study, we identified a key role for LanCL proteins in the dynamic process of post-injury muscle regeneration upon aging. Disruption of all three Lancl genes in mice led to a transient yet significant delay of muscle regeneration in middle-aged but not young adult mice. This impairment does not appear to be due to any change in the quiescent MuSC pool size or its activation. Rather, a delay in the transition from proliferation to differentiation of MuSCs, accompanied by prolonged pro-inflammatory cytokine expression, may have contributed to the observed regeneration phenotype. This aging-dependent phenotype was consistent with the observation that LanCL1 and LanCL2 protein expression were upregulated during regeneration only in the older mice, suggesting that these proteins may have an important function in the aging muscle. Whether the two LanCL proteins play redundant or unique roles is a question that warrants future studies with single KOs. A role of LanCL3 also cannot be definitively ruled out until further investigation.

A key aspect not addressed by our current study is the specific cell type in which LanCL proteins exert their regulatory functions in muscle regeneration during aging. The observed delay in MuSC transition from proliferation to differentiation in LTKO regenerating muscles could reflect a cell-autonomous function of LanCL in the stem cells. However, the prolonged expression of pro-inflammatory cytokines in middled-aged LTKO regenerating muscles would be consistent with a function of LanCLs in immune cells, which impacts MuSC behaviors. A potential involvement of LanCL2 in inflammation has been implicated by its connection to autoimmune conditions and inflammatory diseases (19). The reported role of LANCL2 in ABA-dependent PPARγ activation in macrophages, which suppresses NF-κB/TLR4 signaling and pro-inflammatory cytokine output (41), offers a plausible mechanistic link to the sustained cytokine expression we observed in LTKO muscles. Although mRNA levels of Lancl1 and Lancl2 are relatively low in macrophages throughout the regeneration time course and regardless of age (Fig. 2), information on cell type-specific LanCL protein expression is not available, and a potential LanCL function in macrophages remains a possibility. A third possibility also exists, considering the growing recognition of bi-directional regulation between MuSCs and immune cells (34, 39, 42, 43). While the macrophage is long considered a key contributor to regulating MuSC functions, MuSCs are emerging regulators of immune cells during muscle regeneration. Therefore, it is plausible that LanCL proteins may act in MuSCs to exert immunomodulating functions, which in turn regulates the progression of MuSCs through early stages of regeneration. These possibilities are not mutually exclusive, and future investigations probing the various models may prove fruitful.

An overall well-controlled and timed redox state is another essential component of skeletal muscle regeneration, not only in MuSCs but in all cell types present in the muscle, and this redox balance is progressively destabilized during aging (24, 44). Reactive oxygen species (ROS) generated in injured skeletal muscles impact inflammation and MuSC activation, proliferation, differentiation and self-renewal (24). LanCL1 has been reported to mitigate oxidative stress and preserve mitochondrial function in neurons (5, 9), as well as protecting cancer cells from oxidative stress-induced apoptosis by suppressing JNK signaling (8). Chronic oxidative stress, weaker antioxidant buffering, and altered nitric oxide signaling are all contributing factors to a delayed shift from inflammation to myogenesis during regeneration in aging muscles (25, 44, 45). Glutathione (GSH) is the most abundant small antioxidant molecule in skeletal muscles, and aged MuSCs with lower GSH levels are found to have reduced regenerative capacity (46). LanCL1 and LanCL2 use glutathione as a substrate for post translational C-glutathionylation of various mammalian proteins (12, 47). Structural studies have shown that GSH binds to LanCL1’s active site (18) and LanCL1 has affinity for the signaling protein Eps8, linking LanCL1’s redox regulatory function to cell signaling in the nervous system (48). LanCL1’s antioxidant function is further supported by developmental studies showing that its expression is necessary for neuronal survival under ROS stress (5), and LanCL1 overexpression protects mice with misfolded SOD1G93A from increased oxidative damage (7). It is possible that LanCL deficiency exacerbated the pre-existing redox imbalance in aging muscles due to impaired GSH buffering and compromised antioxidant response, resulting in delayed regeneration.

The function of LanCL proteins in regeneration may also involve their recently discovered role as enzymes that protect the mammalian proteome from accumulation of dehydroamino acid (DHAA) containing proteins. DHAAs have been well documented to accumulate in aging tissues, causing loss of protein function, irreversible cross-linking and insolubility (12, 49, 50). Eliminating damaged proteins is fundamental to maintaining a healthy proteome in aging, without which cellular signaling as well as enzymatic reactions that are essential for post injury muscle repair and regeneration would not proceed normally. DHAA has mainly been reported in the aging lens and brain associated with Alzheimer’s disease (49, 51), and its prevalence and relevance in skeletal muscle has not been studied, possibly due to low abundance and fast protein turnover. Our findings provide the first evidence for LanCL proteins as regulators of the finely tuned process of muscle regeneration, particularly in the context of aging. Determining whether LanCLs play a critical role in mitigating aging-induced protein damage in the regenerating muscle represents an exciting area for future studies.

### Limitation of the study

A potentially significant limitation of the current study is our reliance on systemic/global knockout of LanCLs. No gross phenotype was observed with the LTKO mice, suggesting that any developmental defect of the knockout was unlikely. Nevertheless, we cannot rule out the possibility that our findings could be the result of systemic alterations in the animals rather than a direct effect of LanCL KO on muscle regeneration. Future studies with tissue-specific manipulation of LanCL are warranted.

## ACKNOWLEDGEMENTS

We thank Drs. Kai Zhang, Huaxun Fan, Austin Cyphersmith, and Reza Rajabi-Toustani for expert assistance with fluorescence microscopy and the animal care facility of UIUC for excellent mouse husbandry.

